# A mouse model of classical trigeminal neuralgia via intradural compression of the trigeminal nerve

**DOI:** 10.1101/2025.11.24.690061

**Authors:** Mostafa W. Abdulrahim, Yanxia Chen, Sumil K. Nair, Qian Xu, Yaowu Zhang, Oishika Das, Ryan Gensler, James Feghali, A Karim Ahmed, Christopher M. Jackson, Judy Huang, Youssef G. Comair, Chetan Bettegowda, Xinzhong Dong, Risheng Xu

## Abstract

**Introduction:** Trigeminal neuralgia (TN) is a debilitating orofacial pain condition that adversely affects quality of life. Although heterogeneous, the most common form of TN is classical TN, characterized by paroxysmal bouts of pain in response to otherwise innocuous stimuli. It is believed that classical TN results from neurovascular compression of the trigeminal nerve. However, the underlying pathophysiology of TN is not well understood, thus limiting the development of targeted therapies. Current animal models lack translational relevance, particularly in their inability to replicate intradural nerve root compression, a core anatomic component of TN.

**Methods:** We developed a TN mouse model that achieves intradural nerve root compression via a retro-orbital approach confirmed by anatomic dissection and magnetic resonance imaging. To assess behavioral outcomes, we measured orofacial pain through facial wiping and interaction with a reward stimulus. Pharmacological responsiveness was tested using carbamazepine administration. Mechanistic studies included calcium imaging of trigeminal ganglia (TG), electrophysiologic recordings to measure resting membrane potential and rheobase, and immunohistochemical analysis of the TG.

**Results:** The model elicited orofacial neuropathic pain, substantiated by increased facial wiping and reduced interaction with a reward stimulus, behaviors that suggest both spontaneous and evoked pain. Carbamazepine attenuated these behaviors, suggesting pharmacologic relevance to current TN treatment. Calcium imaging showed heightened spontaneous activity in the TG, and electrophysiologic recordings revealed an increased resting membrane potential and a reduced rheobase. Finally, immunohistochemical studies showed infiltration of CD45+ cells, demyelination and an increase in CGRP expression in the TG, supporting the presence of neuroinflammation after nerve root compression.

**Conclusion:** These findings show that our approach replicates the anatomy and clinical presentation of classical TN in humans. This model may represent a new and robust platform for future mechanistic studies of TN and subsequent preclinical evaluation of therapies in mice.

## 1. Introduction

Trigeminal neuralgia (TN) is a chronic orofacial neuropathic disorder that affects one or more of the dermatomal branches of the fifth cranial nerve.[1, 2] TN constitutes a high clinical burden for patients, widely cited as one of the most debilitating presentations of orofacial pain.[3, 4] It is a heterogeneous condition, of which the most common presentation is termed classical TN.[2] Patients with classical TN report intermittent, shock-like pain episodes that interfere with daily activities.[4, 5] Currently, few effective therapies are available for the medical management of TN. Only carbamazepine is approved by the Food and Drug Administration (FDA) to treat TN; other off-label options include oxcarbazepine, gabapentin, pregabalin, lamotrigine, and phenytoin.[2, 6, 7] The challenge in identifying and developing therapeutic agents is largely due to the limited understanding of TN mechanisms. Additionally, patients often report varying triggers and treatment responsiveness, which makes effectively titrating medications difficult.[8–10] Developing improved therapies for TN requires a better understanding of mechanisms that drive this disease, thus highlighting the need for accurate preclinical models.[11, 12]

Early preclinical models have relied on infraorbital nerve injury to simulate TN. In these models, ligation or chronic constriction is used to injure the infraorbital nerve.[13, 14] Although this approach does induce pain, it represents a peripheral injury. Clinical intraoperative experience has shown that most classical TN presentations result from an aberrant vascular loop compressing the trigeminal nerve root adjacent to the brainstem.[15] Therefore, many researchers have questioned the translatability of these preclinical models as true representations of clinical disease.[11, 12] These concerns are exacerbated by relatively poor evidence derived from animal models of sciatica, which have shown that nerve root and peripheral nerve injury have distinct anatomic, molecular, and neuropathologic, outcomes.[16]

Therefore, we developed a new procedure to compress the trigeminal nerve root intradurally in mice by inserting a silicone-coated filament through the superior orbital fissure. This approach, which we adapted from a technique previously used in rats, is termed intradural trigeminal nerve compression (idTNC). It replicates the classical type of trigeminal neuralgia wherein compression occurs intradurally adjacent to the brainstem. Here, we present our surgical approach to target this region in a murine model. We also characterize pain behaviors, electrophysiologic alterations, immunohistochemical findings, and treatment responsiveness after surgery.

## 2. Materials and methods

### 2.1. Mice

All experiments were performed in accordance with protocols approved by the Animal Care and Use Committee at the Johns Hopkins University School of Medicine. All mice were housed in the Miller Research Building animal facility and weaned at 3 weeks of age. Mice used for this study were 6- to 8-week-old males and females on the C57BL/6J background purchased from Jackson Laboratory.

### 2.2. Surgery

Each mouse was placed into a clean induction chamber and anesthetized by isoflurane 3% to 4% for induction and 1% to 2% for maintenance. Eyes were lubricated to prevent desiccation. A rectal probe was inserted to monitor body temperature. A heating pad placed beneath the mouse was used to maintain body temperature at 37(. The area between the eyes and the snout was gently scrubbed three times with povidone iodine pads followed by alcohol pads to disinfect the skin. A midline incision was made, and the connective tissue underneath was carefully dissected to expose the periorbital bone. A small, loose piece of bone attached to the periorbital bone next to the nasolacrimal duct was carefully removed. A silicone-coated filament (cat #305867, Doccol) marked at 11 mm was inserted at the site where the loose bone was removed and advanced carefully until the marking on the filament reached the periorbital bone. The remaining filament was cut and removed, the skin was closed with 5.0 vicryl suture, and 0.5% lidocaine was injected around the incision site. One milliliter of normal saline was administered subcutaneously and the mice were given intraperitoneal injection of Baytril (2.5mg/kg) once following surgery. Finally, mice were placed in a heating cage and were given diet gel for recovery. Sham mice underwent the same surgery, except the filament was briefly inserted and then immediately withdrawn.

### 2.3. Pain behavior assays

#### 2.3.1. Spontaneous pain assessment

Mice were brought to the testing room for acclimation 3 hours before test initiation. Then, each mouse was placed in separate testing cages covered by aluminum foil to prevent mouse-to-mouse interaction. The mice were left to habituate in this position for 10 minutes before being video recorded for 1 hour. Videos were analyzed in a blinded manner by using Behavioral Observation Research Interactive Software to record the number and duration of face-wiping episodes. Only unilateral face wipes with the forelimb, which are not part of normal grooming, were counted. Mice were recorded for baseline assessment before surgery and then at postoperative days (PODs) 3, 6, 10, and 14.

#### 2.3.2. Evoked pain assessment

For 3 weeks before the surgery, we trained mice in the orofacial pain assessment device (OPAD) cage to receive a sweet reward by pressing their face against metal bars. The sweet solution was made from condensed milk diluted at a 2:1 ratio (water:condensed milk). After the surgery, the mice were deprived of food overnight for 15 hours before starting the test. Each mouse was tested alone in a cage for 20 minutes. For mechanical allodynia testing, mice needed to press metal bars with sharp spikes; for heat allodynia testing, they needed to press 45°C metal bars; and for cold allodynia testing, they needed to press 7°C metal bars. Mice were free to either lick the reward and withstand the stimulus or withdraw if they found it aversive.[17] The OPAD cages were connected to a computer that used ANY-maze software to record the number of licks, number of contacts with each stimulus, and latency time for the first lick. These tests for mechanical, heat, and cold allodynia were conducted at baseline before surgery and at PODs 7, 9, and 11. One day before baseline testing, the mouse buccal hair was shaved carefully with clippers (cat #CL7300-KIT, Kent Scientific) to avoid damaging the whiskers.

Mechanical allodynia was also assessed using the von Frey test. Mice were acclimated in individual chambers for 30 minutes on the day of testing. Mechanical stimuli were applied using a 1.4-g von Frey filament to the facial skin corresponding to the V1 and V2 territories on both the ipsilateral and contralateral sides of the surgery site. Each side was stimulated twice in an alternating manner, with behavioral scores assigned in a blinded fashion. For each mouse, the ipsilateral and contralateral scores were calculated as the average of the two stimulations. Behavioral responses were scored on a scale of 0-4, where 0 = no response, 1 = detection (slight head movement), 2 = quick withdrawal, 3 = face wiping/rubbing, and 4 = vigorous face wiping or attacking.[18] Testing was performed at baseline prior to surgery, and on PODs 1, 3, 7, 10 and 14.

#### 2.3.3. Nesting test

On POD 12, mice were individually habituated in a cage overnight for 12 hours. Then, each mouse was provided with a new, intact nestlet and left in the same cage for another 12 hours. The quality of the resulting nest was then blindly scored with criteria published by Dorninger et al.[19] A score of 1 was given if the nestlet remained completely untouched. A score of 2 indicated partial shredding with at least half of the nestlet left intact. A score of 3 was assigned when the majority of the nestlet was shredded, but the material was scattered with no distinct nest site. A score of 4 reflected a mostly shredded nestlet that was still identifiable but formed a flat nest. Finally, a score of 5 was given if the nestlet was entirely shredded and arranged into a crater-like, clearly defined nest.

### 2.4. Magnetic resonance imaging

All magnetic resonance images were acquired on a 9.4T Bruker spectrometer (Bruker BioSpin Corp., Billerica, MA) using a 25 mm diameter volume coil. The mice were sedated under 1.5% to 2% isoflurane. Brain images were acquired by T2-weighted fast spin-echo rapid imaging with refocused echo (RARE) sequence (repetition time, 3.1 s; echo time, 10 ms; RARE factor, 8; matrix size, 200 x 200; field of view, 2 × 2 cm; slice thickness, 0.5 mm; in-plane resolution, 100 × 100 μm, slice numbers, 22). Quick coronal images were acquired with one acquisition to confirm the position of the filament. Representative images were acquired with six acquisitions in axial, coronal, and sagittal orientation.

### 2.5. Calcium imaging

For calcium imaging experiments, we used Pirt-Cre;GCaMP6s transgenic mice, which express a genetically encoded calcium indicator in trigeminal ganglion (TG) neurons.[20] On POD 7, mice were deeply anesthetized with isoflurane and decapitated. TGs from both the ipsilateral and contralateral sides relative to the surgically compressed trigeminal nerve root, along with the skull base were rapidly harvested and placed in 32(low-sodium Krebs solution—composed of 95 mM NaCl, 2.5 mM KCl, 26 mM NaHCO_3_, 1.25 mM NaH_2_PO_4_-H_2_O, 6 mM MgCl_2_, 1.5 mM CaCl_2_, 25 mM glucose, and 50 mM sucrose—and was bubbled with 95% O_2_/5% CO_2_ for 45 minutes of recovery. Then TGs were transferred to the imaging recording chamber and continuously perfused at 2 mL/min with 95% O_2_/5% CO_2_–bubbled normal Krebs solution, which contained 125 mM NaCl, 2.5 mM KCl, 26 mM NaHCO_3_, 1.25 mM NaH_2_PO_4_-H_2_O, 1 mM MgCl_2_, 2 mM CaCl_2_, and 25 mM glucose. Spontaneous responses of whole-mount neurons were examined in the ipsilateral and contralateral TGs with the Scientifica two-photon system.

### 2.6. Electrophysiology

We conducted patch-clamp recording from ex vivo whole-mount TG neurons. First, we retrogradely labeled TG neurons innervating the skin of orofacial regions by injecting 10 μL of cholera toxin B subunit (CTB) 488 (0.5 mg/mL) into the skin of ipsilateral cheek orofacial regions. One week after the CTB injection, mice were deeply anesthetized with isoflurane and decapitated. Then we rapidly dissected out the ipsilateral TG with the attached nerve bundles and placed them in ice-cold, low-sodium Krebs solution that contained 1 mM kynurenic acid and was bubbled with 95% O_2_/5% CO_2_. Connective tissues on the surface of the TG were carefully removed with fine forceps. TGs with their nerve bundles were then affixed in a recording chamber by a tissue anchor and submerged in normal Krebs solution that was saturated with 95% O_2_ and 5% CO_2_ at room temperature. TGs were exposed to 0.05% dispase II and 0.05% collagenase 4 in the Krebs solution for 10 minutes before being continuously perfused with Krebs solution alone at 2 mL/min. We identified CTB-labeled neurons under epifluorescence illumination and then performed whole-cell patch-clamp recordings on the CTB-labeled V2 TG neurons under the differential interference contrast-infrared (DIC-IR) microscope. The recording electrode internal solution contained the following: 120 mM K-gluconate, 20 mM KCl, 2 mM MgCl_2_, 0.5 mM EGTA, 2 mM adenosine 5′-triphosphate disodium salt (Na_2_-ATP), 0.5 mM guanosine 5′-triphosphate sodium salt (Na_2_-GTP), and 20 mM HEPES. To determine membrane excitability of the CTB-labeled V2 TG neurons, we applied step currents under the current-clamp configuration to the neurons from 0 to 450 pA with increments of 30 pA per step; the duration of each pulse was 1 second. Neuronal excitability was characterized by action potential (AP) threshold, rheobase, resting membrane potential, and numbers of APs evoked by a two-time rheobase stepwise current.

### 2.7. Immunohistochemical analysis

On POD 10, mice were sacrificed, and both ipsilateral and contralateral TGs were harvested and fixed in 4% paraformaldehyde overnight. TG tissues were then transferred into 30% glucose solution for cryopreservation. Afterward, the fixed tissues were embedded in optimal cutting temperature (OCT) compound. The OCT-embedded tissues were sectioned on a cryostat machine and mounted on glass slides for staining. These slides were heated for 30 minutes, then washed three times with phosphate-buffered saline (PBS) for 5 minutes each. Each section was incubated for 1 hour in 200 μL of blocking solution composed of 5% bovine serum albumin (BSA) and 0.3% Triton X-100 in PBS. Rat anti-mouse CD45 primary antibody (1:1000; Biolegend), Rabbit anti-mouse calcitonin gene-related peptide (CGRP) primary antibody (1:500; BMA Biomedicals) and chicken anti-mouse myelin basic protein (MBP) primary antibody (1:200; Sigma-Aldrich) were diluted in PBS containing 0.3% Triton X-100 and 2% BSA and applied to the tissue sections for incubation. After the primary staining, the sections were incubated with goat anti-rat Alexa Fluor 488 secondary antibody (1:500; Invitrogen), donkey anti-rabbit Alexa Fluor 568 secondary antibody (1:500; Invitrogen), and goat anti-chicken Alexa Fluor 647 secondary antibody (1:500; Invitrogen), also prepared in PBS with 0.3% Triton X-100 and 2% BSA. Slides were then mounted with Fluoromount-G with DAPI (Invitrogen) and imaged under a Zeiss LSM700 confocal microscope.

### 2.8. Drug treatment

We assigned mice into four groups, two of which underwent idTNC surgery and two of which underwent sham surgery. One group of idTNC mice and one group of sham-operated mice were treated with 60 mg/kg carbamazepine (TOCRIS), which was administered intraperitoneally daily for 10 days after the surgery.[21] Carbamazepine was dissolved in 10% dimethyl sulfoxide, 5% polysorbate 80, and 85% normal saline. The remaining two groups of mice were treated with vehicle in parallel with the carbamazepine-treated mice.

### 2.9. Statistical analysis

We performed statistical analyses using GraphPad Prism v.10. Statistical comparisons were made by paired Student t test, Mann-Whitney U test, one-way analysis of variance (ANOVA) with Dunnett’s multiple comparisons test, or two-way ANOVA with Sidak’s multiple comparison test or Holm-Sidak’s multiple comparison test. For all bar charts, bars depict the mean. All error bars represent the standard error of the mean, and *n* represents the number of mice analyzed.

## 3. Results

### 3.1. A retro-orbital approach achieves intradural compression of the trigeminal nerve root

Unlike existing rodent models that target peripheral branches of the trigeminal nerve, such as the widely used chronic constriction injury (CCI) of the infraorbital nerve (**Fig. 1A**), we simulate a central injury by compressing the trigeminal nerve root intradurally.[1, 11, 22] To achieve this, we adapted a technique previously used in rats and introduced a silicone-coated filament through the superior orbital fissure to compress the trigeminal nerve root intradurally (**Fig. 1B, C**).[23] While technically feasible in rats, this method posed significant surgical challenges in mice owing to their smaller size and complex skull base anatomy (**Fig. 1C**). Successful filament placement was confirmed postoperatively by T2-weighted magnetic resonance imaging (MRI) in axial (**Fig. 1D**), coronal (**Fig. 1E**), and sagittal (**Fig. 1F**) planes, demonstrating the filament’s position along the skull base and its approach to the trigeminal root. However, filament optimization was essential, as variations in coating dimensions significantly influenced surgical success rates (**Table 1**). Suboptimal filaments often fail to reach the superior orbital fissure, instead lodging in the orbital connective tissues or deviating into the para-esophageal space.

**Figure 1.**
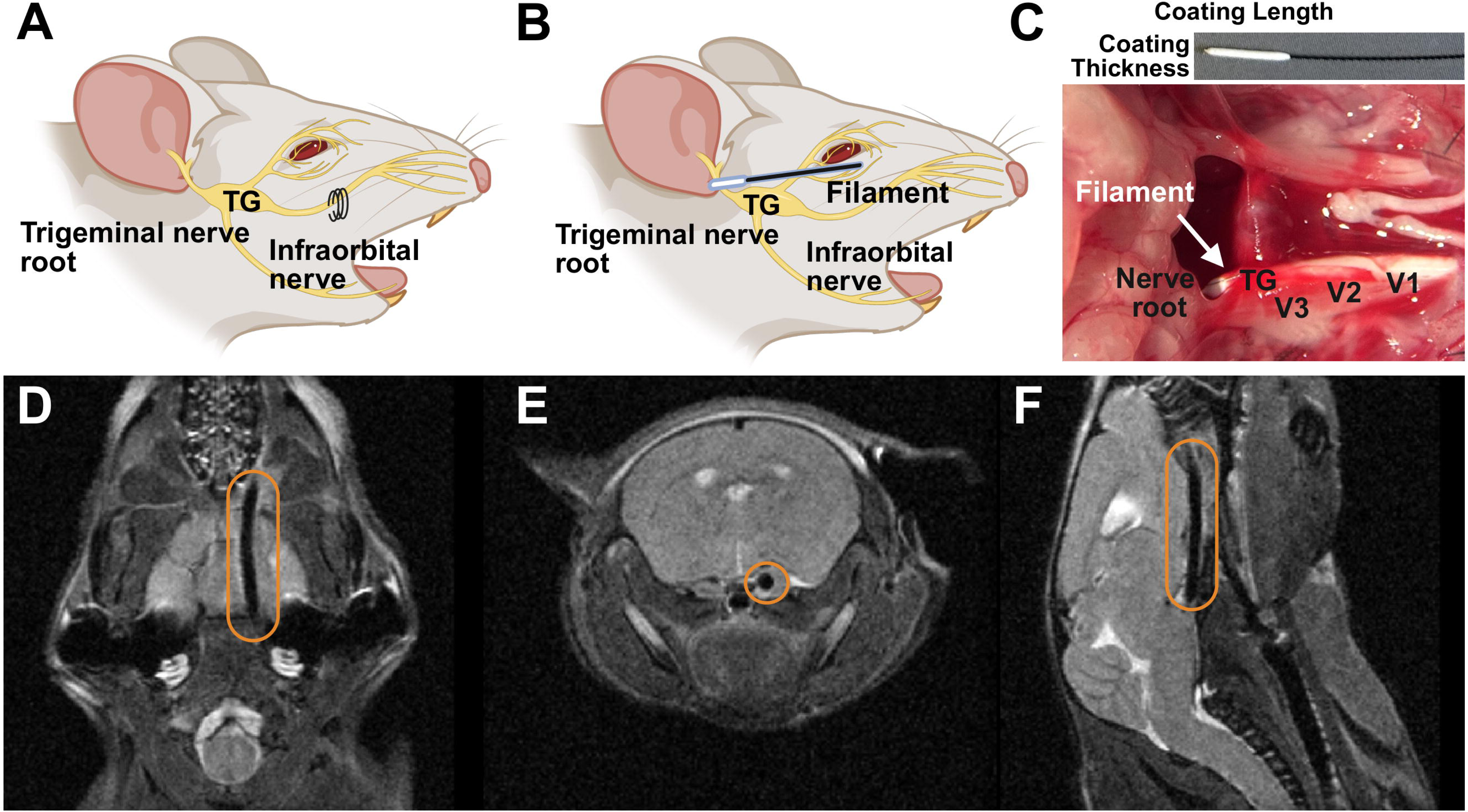
Establishment of the idTNC model in mice. (**A**) Schematic illustration of a peripheral trigeminal nerve injury produced by infraorbital nerve ligation. (**B**) Diagram of the idTNC model, representing intradural nerve root compression of the trigeminal nerve. (**C**) Images of the silicone-coated filament used for compression (top) and an axial view of mouse skull base anatomy showing the filament compressing the trigeminal nerve root intradurally (bottom). (**D–F**) T2-weighted magnetic resonance imaging scans in the axial (**D**), coronal (**E**), and sagittal planes (**F**), confirming intradural compression of the trigeminal nerve. TG: trigeminal ganglion; V1, V2, V3: ophthalmic, maxillary, and mandibular branches of the trigeminal nerve.

**Table 1.** The success rate for different filament coating dimensions.

| Filament coating |  | Performance |  |  |
| --- | --- | --- | --- | --- |
| Length (mm) | Thickness (mm) | Total attempts | Successful attempts | Success rate |
| 3-4 | 0.47 | 8 | 3 | 37.5% |
| 4-5 | 0.47 | 16 | 7 | 43.8% |
| 4-5 | 0.49 | 24 | 13 | 54.2% |
| 4-5 | 0.51 | 10 | 5 | 50.0% |
| **6-7** | **0.58** | **24** | **20** | **83.3%** |

The highest success rate in achieving intradural compression of the trigeminal nerve root was achieved using a filament coated to a length of 6-7 mm and a thickness of 0.58 mm.

### 3.2. The idTNC model results in spontaneous facial pain behavior

We conducted a series of behavioral tests over the course of the experiment to assess postsurgical pain behavior (**Fig. 2A**). First, we investigated the presence of spontaneous, non-evoked pain. Replicating spontaneous pain is crucial because patients with TN often complain of sudden paroxysmal bouts of facial pain without any obvious trigger.[6, 24] Therefore, we measured the spontaneous pain behavior of mice at baseline and several time points after surgery. We observed that after idTNC surgery, both male and female mice had significantly more episodes and longer durations of unilateral face wiping than did sham control mice.[25, 26] This difference began on POD 3, peaked on POD 6 and lasted until POD 10 (**Fig. 2B, C) (Fig. S1A, B**).

**Figure 2.**
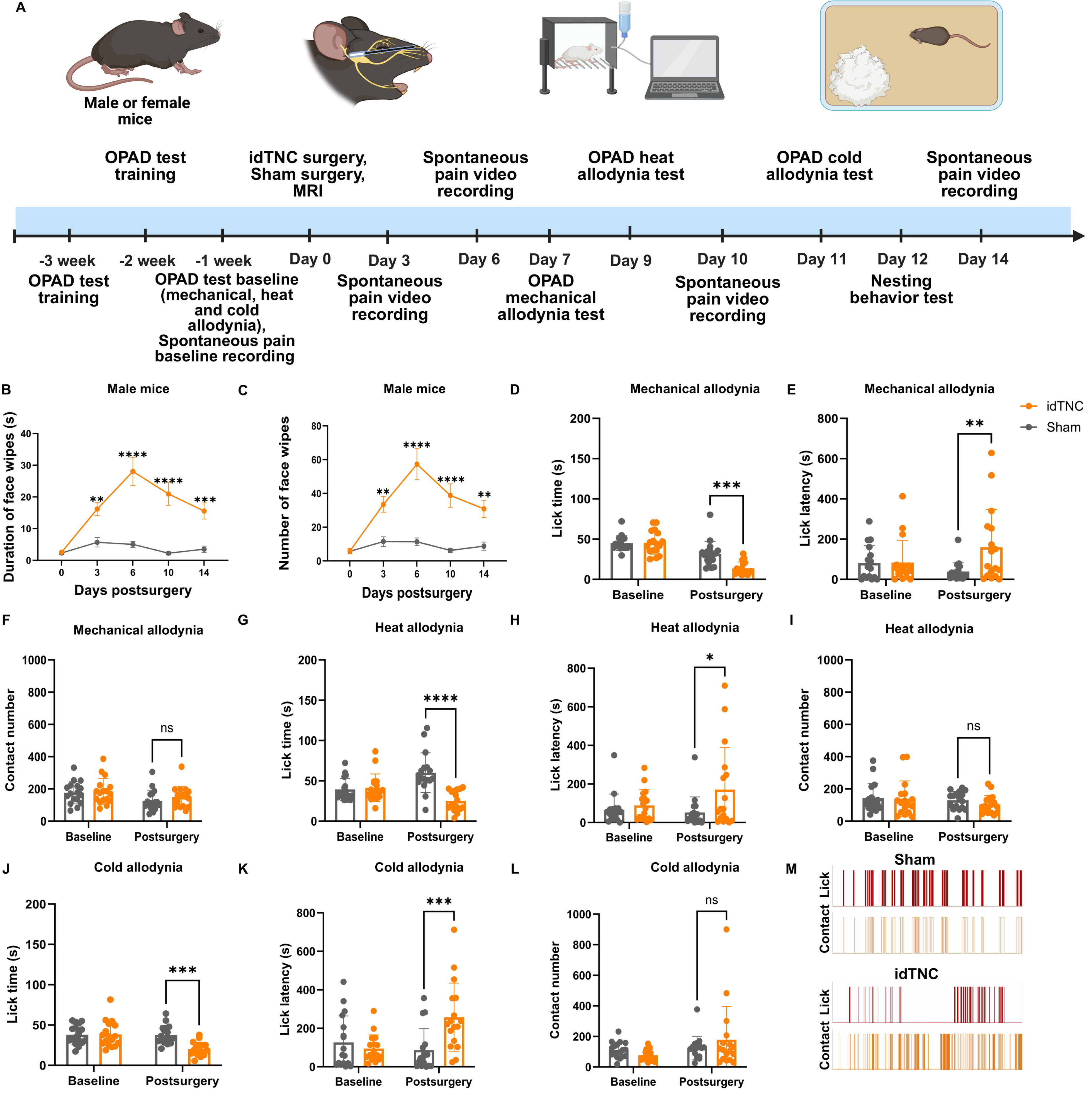
The idTNC model results in spontaneous and evoked facial pain. **(A)** Schematic representation of the study experimental design and timeline. **(B, C)** Male mice that underwent idTNC surgery showed a higher duration **(B)** and number **(C)** of face-wiping episodes compared with sham mice on postoperative days (PODs) 3, 6, 10 and 14. **(D–M)** Evoked pain assessment by OPAD. In the mechanical allodynia test, idTNC mice showed a reduced lick time **(D)** and increased lick latency **(E)** compared with sham-operated mice. **(F)** The number of contacts was comparable between groups in the mechanical allodynia test. **(G, H)** idTNC mice showed significantly reduced lick time and prolonged latency in the heat allodynia test. **(I)** Contact numbers were similar between idTNC and sham-operated mice in the heat allodynia test. **(J, K)** idTNC mice exhibited significantly decreased lick time and increased lick latency in the cold allodynia test. **(L)** Contact numbers were similar between idTNC and sham-operated mice in the cold allodynia test**. (M)** Representative OPAD output diagrams show lick versus contact events for a sham-operated mouse (top) and an idTNC mouse (bottom). (Male mice n=17 for idTNC, n=17 for sham surgery) Statistical analysis: two-way ANOVA with Sidak’s multiple comparisons test. *P<0.05, **P<0.01, ***P<0.001, ****P<0.0001; ns, not significant. OPAD, orofacial pain assessment device.

### 3.3. The idTNC model results in evoked facial pain behaviors

Clinically, patients with TN also report sharp unilateral facial pain that can be triggered by innocuous stimuli such as chewing, light touch, or exposure to cold or heat.[27, 28] Therefore, we used the OPAD test to measure mechanical, heat, and cold allodynia. For male mice on POD 7, mechanical allodynia testing showed that idTNC mice had a significantly shorter lick time of sweet reward (**Fig. 2D**) and a higher lick latency (**Fig. 2E**) compared with sham mice, whereas the number of contacts with the metal spikes was similar between the two groups (**Fig. 2F**). This indicates that while both groups had a similar number of attempts to reach the reward, idTNC mice felt more discomfort in contact with the metal spikes which resulted in increased hesitation and decreased time licking the reward. For female mice the mechanical allodynia testing showed no significant difference in lick time but a higher lick latency in idTNC mice compared to sham mice (**Fig. S1C, D**). Contact number was similar between the two groups (**Fig. S1E**). On POD 9, heat allodynia testing revealed a shorter lick time of sweet reward (**Fig. 2G**) and a higher lick latency (**Fig. 2H**) in male idTNC mice compared with sham mice, while the number of bar contacts remained similar (**Fig. 2I**). Heat allodynia testing in female mice showed a shorter lick time (**Fig. S1F**), but a similar lick latency (**Fig. S1G**) and number of bar contacts (**Fig. S1H**) in idTNC mice compared to sham mice. On POD 11, cold allodynia testing likewise demonstrated shorter lick time (**Fig. 2J**) and higher lick latency (**Fig. 2K**) in male idTNC mice compared to sham mice, with comparable bar contacts between the two groups (**Fig. 2L**). Cold allodynia testing in female mice showed a shorter lick time (**Fig. S1I**), but a similar lick latency (**Fig. S1J**) and number of bar contacts (**Fig. S1K**) in idTNC mice compared to sham mice. Quantitative analysis revealed that idTNC mice made fewer lick attempts than did sham mice when in contact with the noxious stimuli (**Fig. 2M**). Similarly, application of the von Frey filament to the facial skin elicited significantly higher pain score on the ipsilateral side in idTNC mice compared to sham mice (**Fig. S2**). These findings suggest that idTNC may induce mechanical, heat and cold hypersensitivity similar to that experienced by TN patients.

### 3.4. The idTNC model triggers spontaneous activation in the TG

To further investigate the spontaneous pain behavior observed in our study, we directly visualized neuronal activity and spontaneous activities in the ganglia by using Pirt-Cre;GCaMP6s transgenic mice (**Fig. 3A, B**).[29] Calcium imaging showed a significantly higher percentage of spontaneous neuronal firing in TG ipsilateral to the compressed trigeminal nerve root compared to TG from the non-compressed side (**Fig. 3C-E**), further supporting the link between nerve root compression and spontaneous pain behaviors.

**Figure 3.**
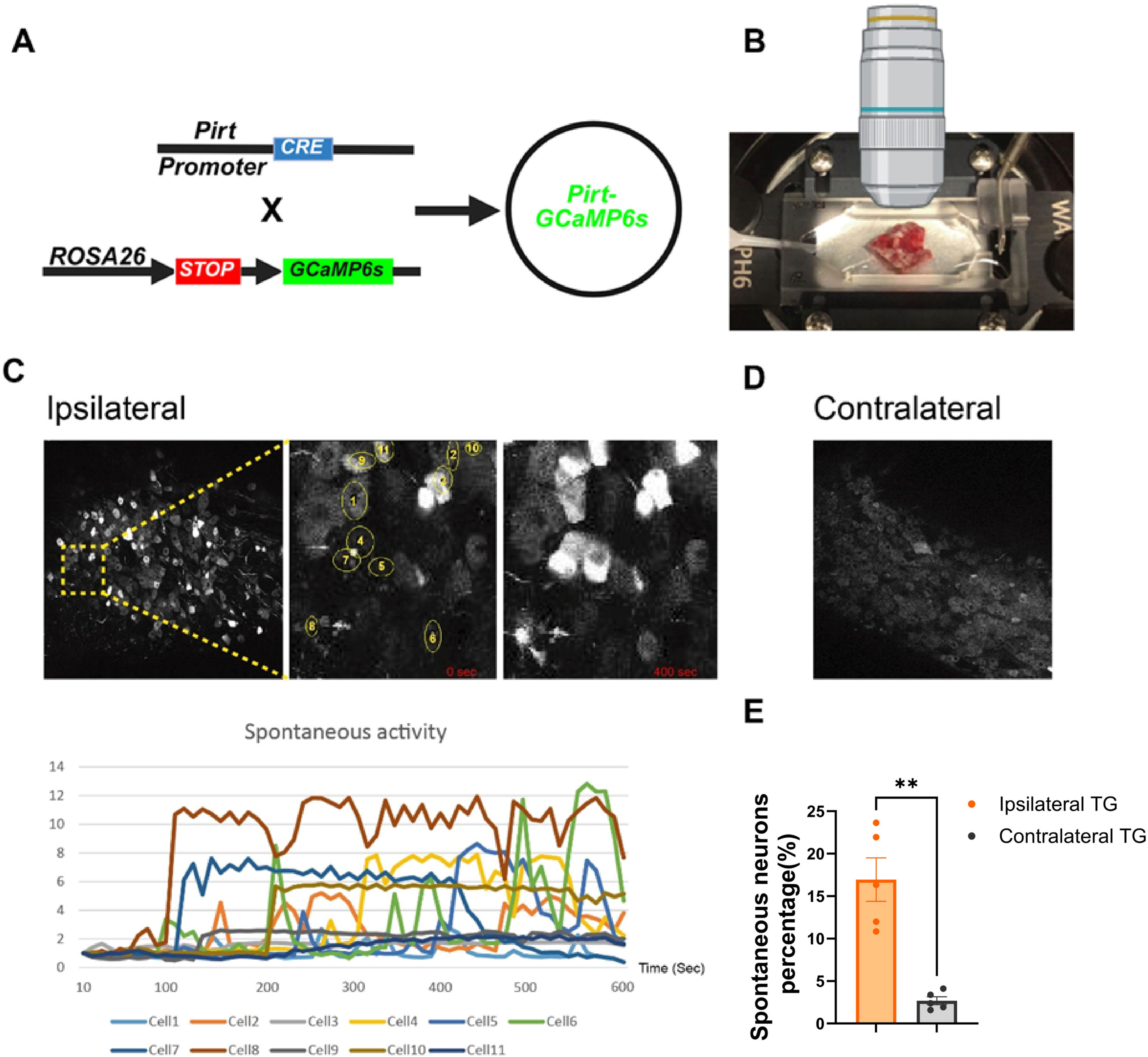
Calcium imaging studies reveal spontaneous neuronal activity in the TG following idTNC surgery. (**A**) Schematic of the Pirt-Cre;GCaMP6s transgenic mouse model used for calcium imaging. Neuronal activation causes fluorescence of the calcium reporter GCaMP6. (**B**) Experimental setup for ex vivo calcium imaging of the TG. (**C**) Representative calcium imaging of TG neurons ipsilateral to idTNC surgery. Left: Wide-field view showing neuronal populations. Middle: Magnified region with numbered spontaneously active neurons circled at the 0 second time point. Right: Changes in fluorescence intensity at 400 seconds. Bottom: Spontaneous neuronal activity over time in TG neurons ipsilateral to idTNC compression. Each colored line represents the activity of an individual neuron. (**D**) Representative calcium imaging of TG neurons contralateral to idTNC surgery. (**E**) Quantification of the percentage of spontaneously active neurons in TGs ipsilateral and contralateral to idTNC surgery. Statistical analysis: paired Student t test. \*\**P*<0.01 (*n*=5 ipsilateral TG, *n*=5 contralateral TG).

### 3.5. The idTNC model causes hyperexcitability in the trigeminal nerve

We conducted electrophysiologic studies to evaluate the neuronal activity and plasticity of small V2 TG neurons innervating orofacial cutaneous regions after idTNC surgery (**Fig. 4A-C**). The idTNC model induced hyperexcitability in these neurons, accompanied by significant changes in their electrophysiologic properties. Specifically, resting membrane potentials were significantly increased in the idTNC group compared with those in the sham-operated animals (**Fig. 4D**). This depolarization reflects a fundamental alteration in the neuronal state, contributing to heightened excitability.[5] Additionally, the rheobase, the minimum current needed to evoke an AP, was also significantly decreased in TG neurons after idTNC surgery, suggesting increased sensitivity to incoming stimuli (**Fig. 4E**).[30] Moreover, idTNC TG neurons exhibited lower AP amplitudes and longer AP durations than did neurons from sham-operated mice (**Fig. 4F, G**). This shift indicates a profound alteration in neuronal firing behavior, further underscoring the hyperexcitability induced by the idTNC model.

**Figure 4.**
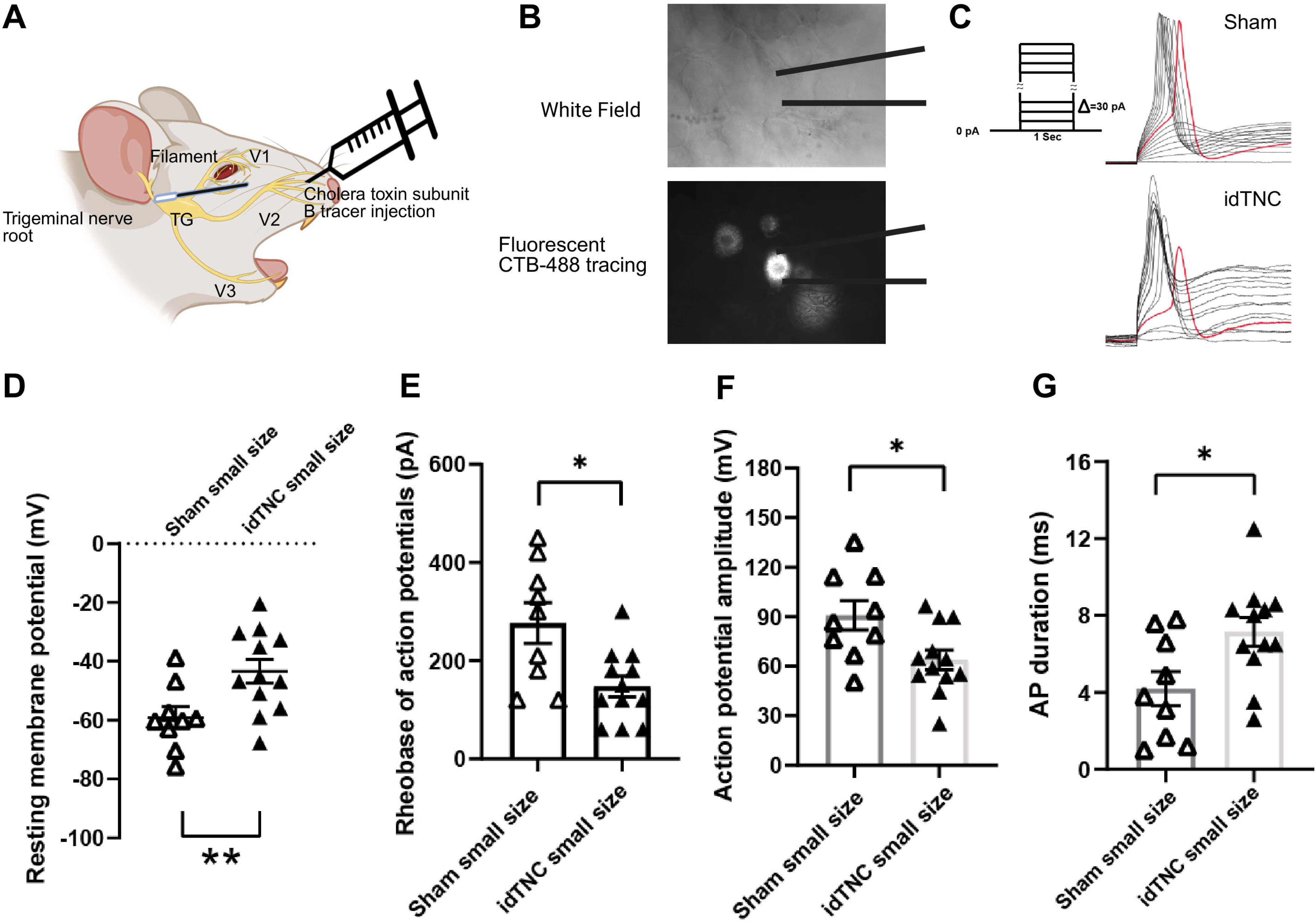
Electrophysiology studies demonstrate a hyperexcitable state of the trigeminal nerve after idTNC surgery. (**A**) Schematic representation of patch-clamp experiments. (**B**) Whole mount of TG recording under cholera toxin subunit B (CTB) tracing from V2 branch. (**C**) Sample action potential traces recorded from a small CTB-labeled V2 TG neuron in sham-operated and idTNC mice with protocol for action potential recording. (**D**) Small neurons from mice that underwent idTNC surgery showed a significantly higher resting membrane potential than those from sham-operated mice. (**E**) The action potential rheobase was significantly lower in small neurons from idTNC mice than in those from sham-operated mice. (**F**) Action potential amplitude was significantly lower in small neurons from idTNC mice than in those from sham-operated mice. (**G**) Action potential duration was significantly greater in small neurons from idTNC mice than in those from sham-operated mice. (**D-G**) Statistical analysis: Mann-Whitney U test. \**P*<0.05, \*\**P*<0.01 (*n*=12 idTNC, *n*=9 sham). AP; action potential; TG: trigeminal ganglion; V1, V2, V3: ophthalmic, maxillary, and mandibular branches of the trigeminal nerve.

### 3.6. The idTNC model results in immune cells migration, demyelination and an increase in CGRP expression in the trigeminal ganglia

We next examined neuroinflammation, demyelination and CGRP expression in the TG after idTNC. We stained TGs for CD45 to identify immune cell infiltration on compressed (ipsilateral) and non-compressed (contralateral) sides (**Fig. 5A**). This showed a significantly higher expression of CD45+ cells in the ganglia ipsilateral to surgical nerve root compression, indicating increased immune cell infiltration in response to nerve root injury (**Fig. 5B**).[31, 32] MBP staining showed a lower average intensity per region of interest (ROI) in the TG on the compressed side compared to the non-compressed side, indicating the presence of increased demyelination following idTNC surgery (**Fig. 5C-D**).[33] Moreover, CGRP, a neuropeptide released from sensory neurons after injury that contributes to pain sensitization and downstream nociceptive signaling, was expressed at a higher level in the ipsilateral TG compared to the contralateral side (**Fig. 5E-F**).[34]

**Figure 5.**
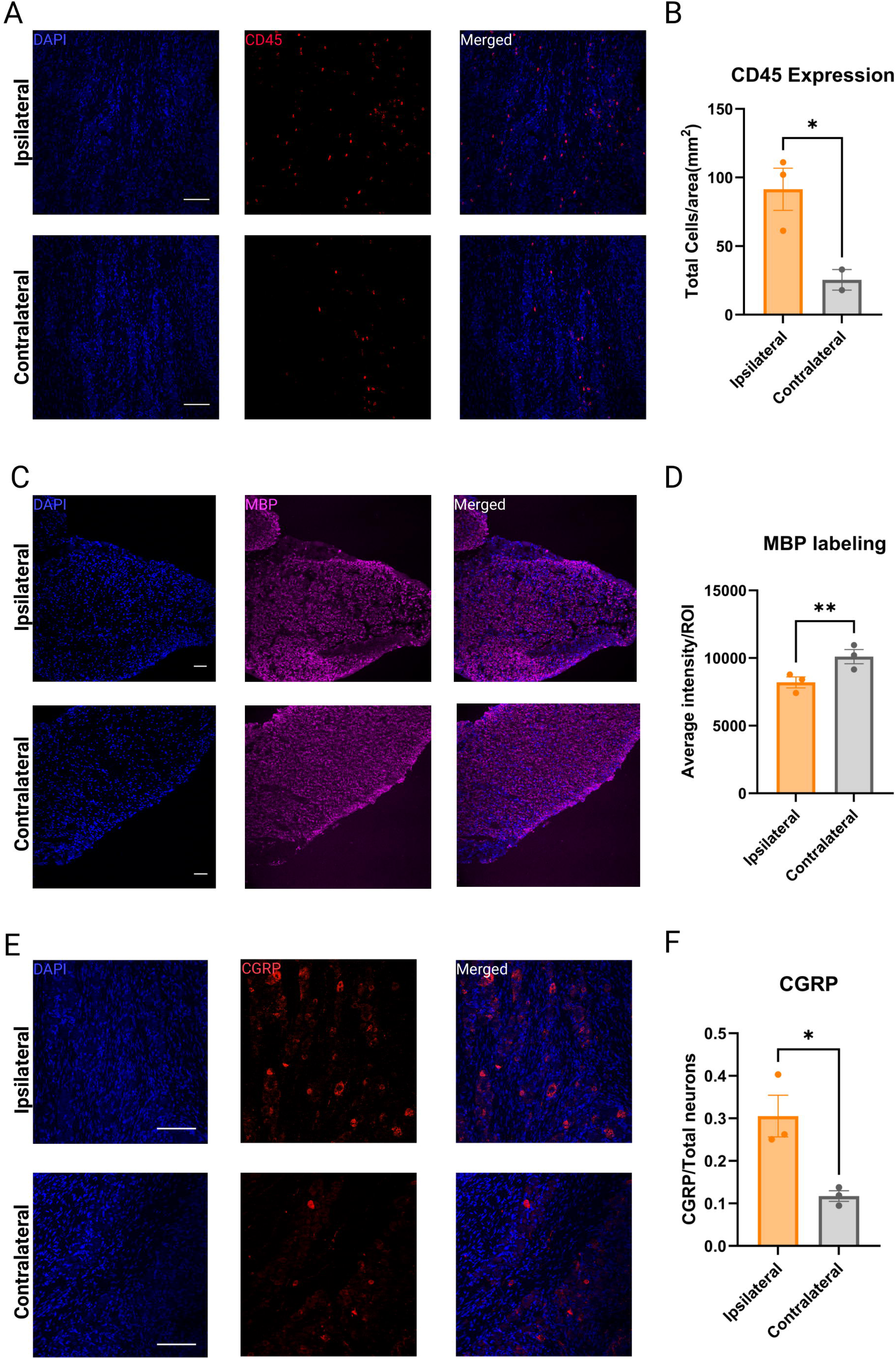
Immunohistochemistry studies reveal increased CD45+ cellular infiltration, demyelination, and higher expression of CGRP in the TG after idTNC surgery. (**A**) TG ipsilateral to the idTNC surgery (top) exhibits significantly higher expression of CD45+ cells compared with TG contralateral to idTNC surgery (bottom). (**B)** Quantification showed a significantly higher number of CD45+ cells in TG on the side ipsilateral to idTNC surgery than in TG on the contralateral side. (**C-D**) Ipsilateral TG (top) exhibits significantly lower MBP average intensity per ROI compared with contralateral TG (bottom). (**E-F**) Ipsilateral TG (top) exhibits significantly higher expression of CGRP normalized to the total number of neurons, compared with contralateral TG (bottom). The data represent average values from 3 ipsilateral TGs for CD45, MBP and CGRP staining, and from 3 contralateral TGs for MBP and CGRP, as well as 2 contralateral TGs for CD45. For each ganglion 9 images from 3 slide sections were captured. Statistical analysis: paired Student t test. \**P*<0.05, \*\**P*<0.01.

### 3.7. Carbamazepine attenuates pain behaviors in mice after idTNC

After establishing the presence of spontaneous and evoked pain in mice after idTNC, we assessed the effect of carbamazepine on pain behavior.[7, 10] Male and female carbamazepine-treated idTNC mice exhibited a significant decrease in face-wiping episode duration compared with vehicle-treated idTNC mice after 6 days of injections, suggesting a decrease in spontaneous pain (**Fig. 6A, B**). The idTNC mice also exhibited greater lick time with carbamazepine treatment than with vehicle treatment in the OPAD tests for mechanical and cold allodynia (**Fig. 6C-K**). We also tested mouse nesting behavior, an established indicator of mouse welfare that can be impaired by pain and discomfort.[35, 36] Nesting scores were significantly lower in vehicle-treated idTNC mice than in carbamazepine-treated idTNC mice (**Fig. 6L-P**), indicating mouse responsiveness to carbamazepine treatment.[19]

**Figure 6.**
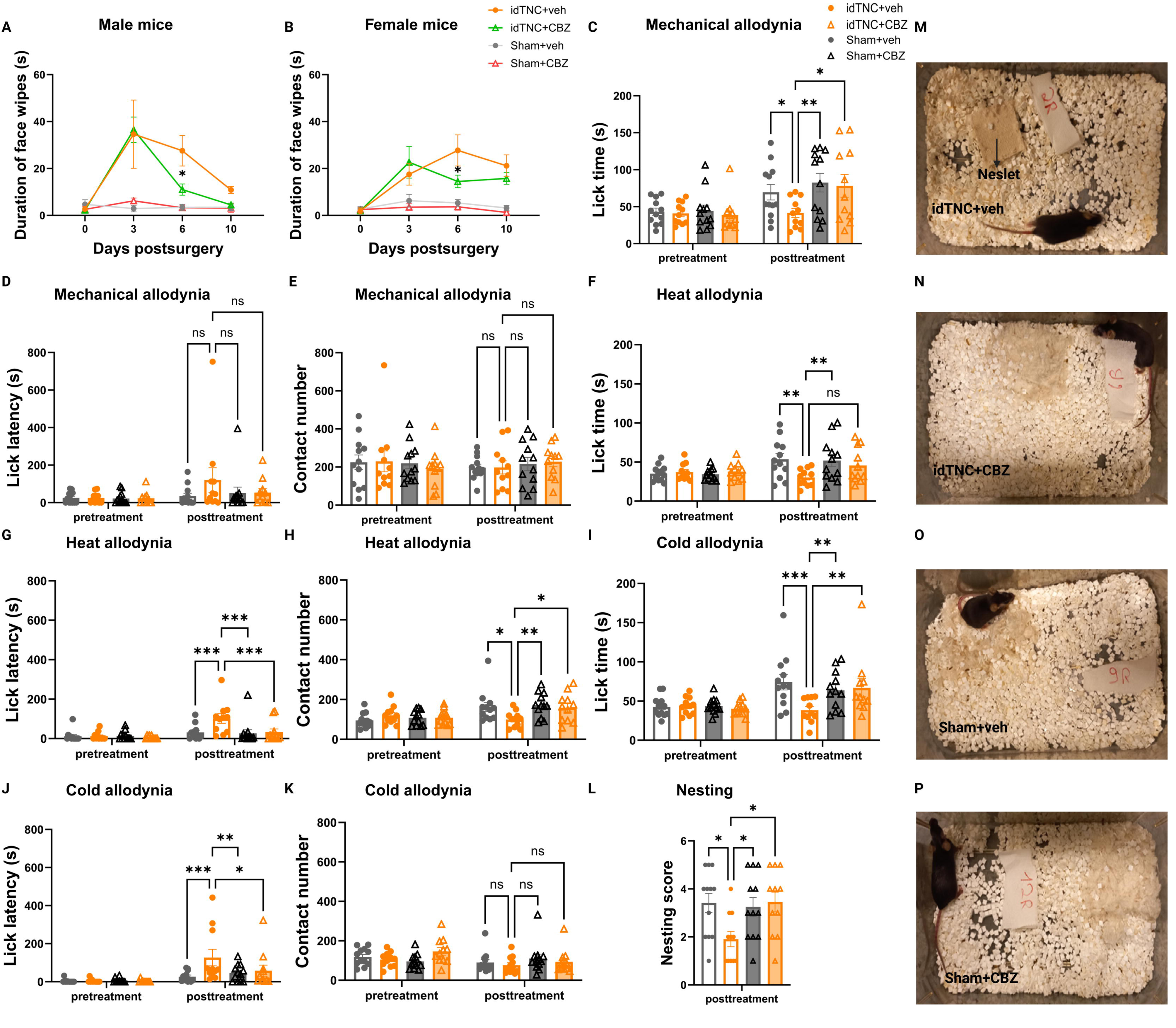
Carbamazepine (CBZ) treatment attenuates spontaneous and evoked facial pain after idTNC. (**A, B**) Spontaneous pain behavior: On postoperative day 6, face wiping episodes in both male (**A**) and female (**B**) idTNC mice were shorter in duration after CBZ treatment than after vehicle (veh) treatment. (**C-K**) OPAD testing for mechanical, cold, and heat allodynia with combined male and female data. (**C**) In the mechanical allodynia test, vehicle-treated idTNC mice showed significantly less lick time than did CBZ-treated idTNC mice. (**D**) No significant difference was observed for the lick latency between the different groups in the mechanical allodynia test. (**E**) Contact number was similar between CBZ-treated idTNC and vehicle-treated idTNC mice in the mechanical allodynia test. (**F**) In the heat allodynia test, vehicle-treated and CBZ-treated idTNC mice exhibited no significant difference in lick time. Vehicle-treated idTNC mice exhibited significantly increased lick latency (**G**) and decreased contact number (**H**) compared with CBZ-treated mice in the heat allodynia test. In the cold allodynia test, vehicle-treated idTNC mice exhibited significantly less lick time (**I**) and greater lick latency (**J**) than did CBZ-treated idTNC mice. (**K**) Contact number was consistent between groups in the cold allodynia test. (**L**) Vehicle-treated idTNC mice had a significantly lower nesting score than did CBZ-treated idTNC mice. (**M–P**) Representative images of nesting behavior: (**M**) vehicle-treated idTNC mouse (idTNC+veh), (**N**) CBZ-treated idTNC mouse (idTNC+CBZ), (**O**) vehicle-treated sham-operated mouse (Sham+veh), (**P**) CBZ-treated sham-operated mouse (Sham+CBZ). (Female mice: *n*=5 for idTNC+veh, *n*=6 for Sham+veh, *n*=6 for idTNC+CBZ, *n*=6 for Sham+CBZ; male mice: *n*=6 for idTNC+veh, *n*=6 for Sham+veh, *n*=5 for idTNC+CBZ, *n*=6 for Sham+CBZ) Statistical analysis: (**A, B**) two-way ANOVA with Sidak’s multiple comparisons test; (**C-K**) two-way ANOVA with Holm-Sidak’s multiple comparisons test; (**L**) one-way ANOVA with Dunnett’s multiple comparisons test. \**P*<0.05, \*\**P*<0.01, \*\*\**P*<0.001; ns, not significant.

## 4. Discussion

Classical TN pain is thought to develop from a neurovascular compression of the trigeminal nerve root,[28] but how this injury leads to pain phenotypes is less understood mechanistically. Improving pharmacotherapeutics necessitates a more thorough understanding of TN pathophysiology, thus highlighting the importance of accurate preclinical models.[11] Most current TN models have significant limitations. For example, the CCI model involves ligation of the infraorbital nerve. Although this model produces an orofacial pain phenotype, the approach represents a peripheral injury, rather than a central nerve root injury.[22, 37] Recently, Ding et al.[38] developed the foramen lacerum impingement (FLIT) model, in which nerve root compression is achieved through the foramen lacerum. However, this compression is extradural, rather than intradural. Luo et al.[23] successfully implemented intradural compression of the trigeminal ganglion in rats, and establishing a similar model in mice would offer advantages for investigating TN. Mice are not only more cost-effective to maintain than rats, but they are also more routinely used to test genetic alterations.[39] Therefore, we sought to develop a robust and reliable mouse model of classical TN. Our surgical approach achieved intradural, trigeminal nerve root injury that may more accurately reflect classical TN than earlier models.

TN is a complex orofacial pain disorder, characterized by both spontaneous and evoked pain symptoms. Not only do patients report hypersensitivity to normally innocuous stimuli such as light touch, they also experience intermittent episodes of severe pain while at rest.[10, 40, 41] These two types of pain are mechanistically distinct and associated with different patterns. Paroxysmal pain that occurs in the absence of an external stimulus is hypothesized to result from spontaneous discharge within the ganglia, whereas allodynia is thought to arise in part from hypersensitization of the peripheral nerve itself.[42, 43] In our model, we successfully achieved both spontaneous and evoked pain behaviors after intradural nerve root injury in both male and female mice. We note that male mice experience more orofacial pain than female mice in our model. Our experimental paradigm utilized female mice without synchronization of the estrous cycle; estrous stage was not monitored at the time of behavioral testing. Since ovarian hormone fluctuations can influence nociceptive thresholds and neuropathic pain mechanisms, we note this as a potential biological contributor to the sex difference observed in our results.[44]

Historically, researchers have evaluated evoked pain behaviors in mouse models of TN with the von Frey test. In that test, an investigator applies monofilaments of increasing force in series to the animal to determine at what threshold a stimulus leads to a withdrawal response.[37] However, interpretation of this approach is limited because stimuli are forcibly applied. In our study, we mainly assessed evoked pain through an OPAD, in which mice are trained to obtain a reward. While obtaining the reward, they are exposed to mechanical, cold, or hot stimuli. In this test, mice voluntarily interact with the OPAD, allowing the device to capture more complex, decision-based behaviors that reflect the animals’ pain perception.[17, 26] Interestingly, although idTNC and sham-operated mice made a similar number of contacts with the stimulus, mice in the idTNC group exhibited reduced licking behavior. This deliberate avoidance of painful stimuli may represent altered pain processing and mimic the behavior of patients with TN who often avoid activities such as chewing or eating that can provoke pain episodes.[45]

When using preclinical models of TN, it is also essential to demonstrate that pain behaviors are amenable to carbamazepine treatment to accurately reflect the clinical course. For example, the study by Ding et al.[38] that used the FLIT model was limited because it did not assess the efficacy of carbamazepine treatment in reversing pain behaviors. In our study, the orofacial pain behaviors produced by our model were modifiable with carbamazepine treatment, further supporting the validity of our approach.

Given that patients with TN experience both spontaneous pain and heightened sensitivity to innocuous stimuli, researchers have suggested that nerve root compression leads to increased discharge within the TG. This phenomenon results in spontaneous firing within the ganglia, causing neurons to become overly responsive and to produce a burst of activity in response to any trigger.[5, 10, 27] To query whether our TN model similarly resulted in aberrantly active ganglia, we performed calcium imaging of TG after idTNC surgery. We observed increased spontaneous neuronal activity in the TG on the compressed side compared with that on the non-compressed side, which may explain the paroxysmal pain behavior observed in mice. Moreover, the allodynia observed in this model may be attributed to electrophysiologic changes in small-diameter V2 trigeminal neurons. Postsurgical recordings showed a more depolarized resting membrane potential and a lower rheobase in idTNC mice than in sham mice. These alterations suggest a hyperexcitable neuronal state and peripheral sensitization of the nerve, conditions that align with the evoked pain behaviors observed in our model.[46]

Although physical compression of the trigeminal nerve root is thought to be the primary driver of disease pathology in classical TN, very little is known about molecular downstream mechanisms that contribute to this disease. Recent studies in patients have revealed that inflammatory processes may play a pivotal role in TN and could represent new therapeutic targets. When Ostertag et al.[32] analyzed cerebrospinal fluid (CSF) samples from patients with TN and healthy controls, they found elevated levels of proinflammatory cytokines (IL-9, IL-18, and IL-33), chemokines (RANTES and ENA-78), the tumor necrosis factor superfamily (TRAIL and sCD40L), and growth factors (EGF, PDGF-AB/BB, and FGF-2). A 2019 study by Ericson et al.[47] revealed that TRAIL and TNF-β were elevated in the CSF of patients with TN prior to microvascular decompression. After surgery, these biomarkers decreased and correlated with clinical pain relief. Furthermore, studies of trigeminal specimens from patients undergoing microvascular decompression with rhizotomy have demonstrated demyelination in the trigeminal nerve root. This finding has been proposed as a key pathological mechanism in TN, contributing to ectopic activity and abnormal pain signaling.[48] Additionally, a clinical study detected elevated CGRP levels in both CSF and peripheral blood of TN patients, highlighting a potential role for this neuropeptide in pain sensitization and disease pathophysiology.[49] To investigate inflammation and pathological changes in the TG following filament insertion, we immunohistochemically stained TG and found that the ganglia on the compressed side exhibited higher levels of CD45+ cells, more demyelination, and a higher expression of CGRP than did the ganglia on the non-compressed side. The increase in CD45+ cells highlights a localized inflammatory response triggered by filament insertion, which may reflect the inflammatory changes observed in TN patients. Likewise, the demyelination we observed parallels findings in human surgical specimens, where disruption of myelin was suggested to drive ectopic activity and pathological nociceptive signaling. Finally, the upregulation of CGRP expression in our model mirrors prior clinical studies reporting elevated CGRP levels in TN patients, further supporting the relevance of our model in recapitulating TN disease mechanisms.

Despite the improved internal validity of our model relative to the popular CCI model, we acknowledge that TN is a heterogeneous condition and that our model only reflects classical TN pain resulting from neurovascular compression. These findings should not be used to draw conclusions regarding atypical or multiple sclerosis–related TN pain, which represent distinct pathophysiologic mechanisms.[50, 51] However, we believe that this model is the first to replicate intradural, compression of trigeminal nerve in mice. This model offers a clinically relevant platform for investigating the mechanisms of TN and other neuropathic pain conditions. Future studies incorporating detailed cellular and molecular analyses of the TG and surrounding tissue may further elucidate the inflammatory and neuroplastic changes that drive TN pathology. Advancing our understanding of TN pathophysiology is vital for the development of targeted, long-lasting therapeutic strategies for this patient population.

## Abbreviations

TN: Trigeminal neuralgia
TG: Trigeminal ganglia
idTNC: intradural trigeminal nerve compression
POD: Postoperative day
OPAD: Orofacial pain assessment device
AP: Action potential
CGRP: Calcitonin gene-related peptide
MBP: Myelin basic protein
CBZ: Carbamazepine
CSF: Cerebrospinal fluid

## Acknowledgments

Illustrations/diagrams were created with BioRender and GraphPad Prism v.10. We thank Santosh K. Yadav, and Marie-France Penet from the MRB Molecular Imaging Service Center at Johns Hopkins University for their help with rodent MRI sequences.

## Author contributions

R.X., M.W.A., and Y.C. conceptualized the study. M.W.A., Y.C., S.K.N., Q.X., Y.Z., and O.D. performed experiments and/or analyzed data. R.X., M.W.A, Y.C., S.K.N., Q.X., R.G., J.F., AK.A C.M.J., C.B., J.H., Y.G.C., and X.D. reviewed data. M.W.A, R.X., and S.K.N. wrote the manuscript, and all authors edited and approved the manuscript.

## Data availability

All data are available upon request from the corresponding author.

## Declarations

**Figure S1.**
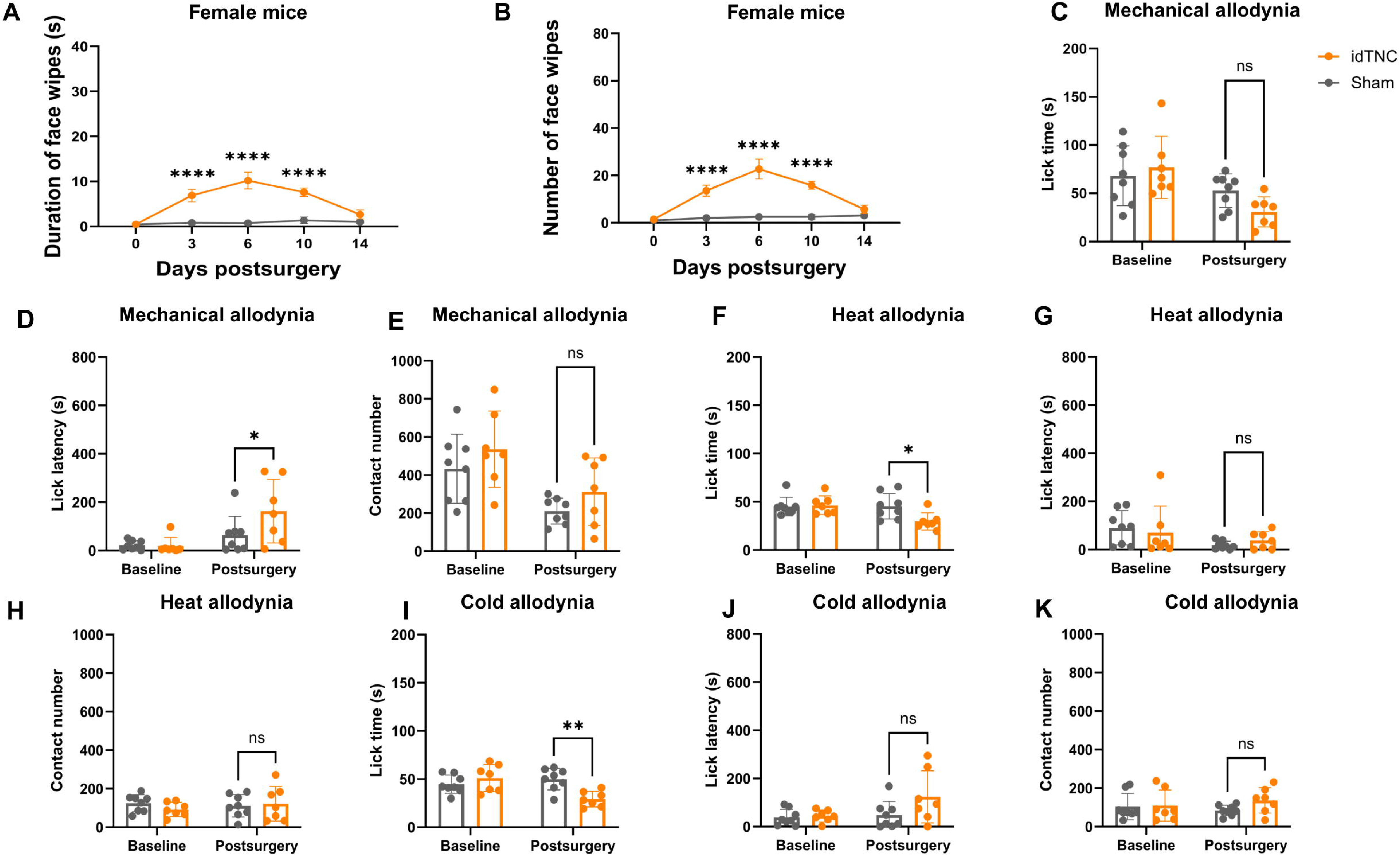
The idTNC model results in spontaneous and evoked facial pain in female mice. (**A, B**) female mice that underwent idTNC surgery showed a higher duration (**A**) and number (**B**) of face-wiping episodes compared with sham mice on postoperative days (PODs) 3, 6, and 10. (**C–K**) Evoked pain assessment by OPAD. In the mechanical allodynia test, idTNC mice showed no significant difference in lick time (**C**) but a significant increase lick latency (**D**) compared with sham-operated mice. (**E**) The number of contacts was comparable between groups in the mechanical allodynia test. (**F, G**) idTNC mice showed significantly reduced lick time but no difference in lick latency in the heat allodynia test. (**H**) Contact numbers were similar between idTNC and sham-operated mice in the heat allodynia test. (**I, J**) idTNC mice exhibited significantly decreased lick time and a similar lick latency in the cold allodynia test. (**K**) Contact numbers were similar between idTNC and sham-operated mice in the cold allodynia test. (Female mice *n*=7 for idTNC, *n*=8 for sham surgery). Statistical analysis: two-way ANOVA with Sidak’s multiple comparisons test. \**P*<0.05, \*\**P*<0.01, \*\*\*\**P*<0.0001; ns, not significant. OPAD, orofacial pain assessment device.

**Figure S2.**
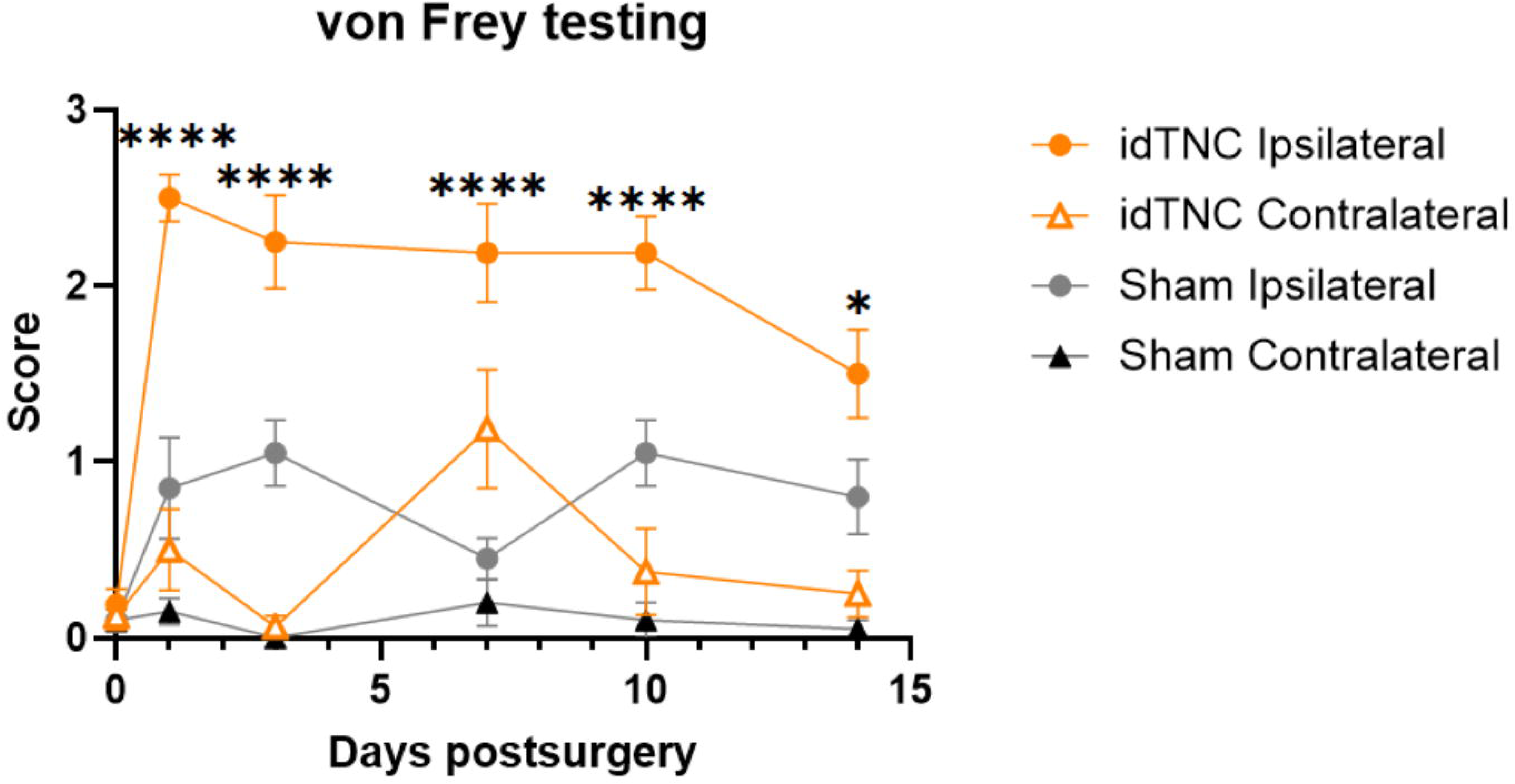
Mechanical allodynia measured by orofacial von Frey test. On post operative days 1, 3, 7, 10 and 14 idTNC mice exhibited significantly higher mechanical sensitivity on the ipsilateral side compared with sham mice. Data are shown as mean values with ± SEM. (*n*=8 for idTNC, *n*=10 for sham surgery) Statistical analysis: two-way ANOVA with Tukey’s multiple comparisons test. \**P*<0.05, \*\*\*\**P*<0.0001.

